# Germline Blastomeres transcriptomics in the presence or absence of PIE-1 during *C. elegans* early embryogenesis

**DOI:** 10.64898/2026.02.14.705883

**Authors:** Pauline Ponsard, François-Xavier Stubbe, Pauline Tricquet, Craig C. Mello, Damien Hermand

**Author notes:** Department of Molecular and Cellular Biology, Faculty of Sciences, University of Geneva, Geneva, Switzerland (François-Xavier Stubbe). Department of Microbiology, Tumor and Cell Biology, Karolinska Institute, Solna 171 76, Sweden (Pauline Tricquet). These authors contributed equally to this work (Pauline Ponsard, François-Xavier Stubbe).

## Abstract

In *Caenorhabditis elegans (C. elegans)* embryos, specification of the germ lineage relies on the maternal protein PIE-1. Here, we isolated somatic and germline blastomeres and performed specific transcriptomics in the presence or absence of PIE-1. These experiments enforce that PIE-1 maintains the identity of germline blastomeres by downregulating the accumulation of somatic RNAs but they also reveal a role for PIE-1 in sustaining the steady-state level of both maternally contributed as well as *de novo* transcribed germline-specific mRNAs in the early embryo. The present dataset represents a first blastomere-specific transcriptomic analyses of PIE-1 role and will serve as a comprehensive resource to highlight the mechanistic details of PIE-1 function in defining germline blastomere transcriptional identity.

## Background & Summary

Safeguarding germ cell identity from somatic differentiation during embryogenesis is of critical importance. This is why germline blastomeres (early embryonic cells) are segregated from somatic blastomeres at the very beginning of embryogenesis and many biological processes are distinctly regulated in these two cell types. In many organisms, germline blastomeres appear to enter a transcriptionally quiescent state during early development, which is proposed to protect them from adopting a somatic fate ^1^. Typically, in *C. elegans*, Pol II-mediated transcription is activated in somatic blastomeres at the 4-cell stage but the germline blastomere remain transcriptionally quiescent until the 100-cell stage ^2,3^.

In *C. elegans*, Pol II-mediated transcription is proposed to be globally inhibited in the germline blastomeres by the PIE-1 protein ^3^ although this has not been demonstrated at single gene resolution. PIE-1 is a CCCH-type zinc finger protein characterized by two zinc finger domains (ZF1 and ZF2) separated by an arginine and serine-rich domain (RS-rich domain) ^2^. PIE-1 is present in the oocytes and is maternally deposited in the one-cell embryo ^3^. It segregates exclusively to the early germline lineage consisting of P1-P2-P3-P4 cells, where it accumulates in the nucleus, consistent with its role as a transcriptional inhibitor ^3,4^.

PIE-1 is degraded around the 100-cell stage in the two P4 daughter cells, Z2 and Z3, coinciding with a transient activation of transcription ^2^. In the absence of PIE-1, both germline and somatic blastomeres initiate transcription simultaneously, and the germline blastomere P2 adopts a fate similar to its sister somatic cells, leading to embryos with an excess of pharyngeal and intestinal cells (PIE stands for pharyngeal and intestinal cells in excess), ultimately resulting in embryonic lethality ^5^.

In this study, we present a transcriptomic dataset supporting that PIE-1 is crucial for maintaining the identity of germline blastomeres by sustaining the steady-state level of maternally contributed mRNAs. In addition, when PIE-1 is absent, somatic RNAs accumulate in germline blastomeres, further compromising their germline identity. These data pave the way for deciphering the molecular mechanisms underlying the function of PIE-1 in defining germline blastomeres transcriptional identity.

## Methods

### Germline and somatic blastomeres have distinct transcriptomes

The specificity of the transcriptomic landscape of germline and somatic blastomeres is a long-standing question in developmental biology. To address this question, we aimed to sort out the *C. elegans* germline blastomeres (referred to as P2, P3, P4, Z2, Z3 along early embryonic divisions) from the somatic blastomeres. We first set a cell sorting assay during early embryogenesis. It is not trivial to identify markers that strictly localize in germline blastomeres at all developmental stages. Based on literature mining and trials, we selected the Argonaute protein WAGO-4 ^6^.

Using embryo dissociation and FACS sorting on a *wago-4::gfp* (Seroussi et al., 2023) strain, we separated GFP-positive (GFP+) from GFP-negative (GFP-) blastomeres **(Figure 1)**, which were then subjected to bulk RNA-seq. The resulting transcriptomic landscapes showed a clear distinction between GFP+ and GFP- blastomeres, as demonstrated by a Principal Component Analysis (PCA) where 91% of the variance was attributable to the differences between GFP+ and GFP- blastomeres **(Figure 2A)**. These two types of sorted blastomeres exhibited dramatically distinct transcriptomic landscapes. Overall, 6056 transcripts, such as *lpr-3,* were enriched in GFP-blastomeres. Conversely, 3703 transcripts, including *pie-1*, were enriched in GFP+ blastomeres (Fold change > 2 and adjusted *p* value < 0.01). Finally, among the differentially expressed transcripts (adjusted *p* < 0.01), 1272 had an absolute fold change below the threshold and therefore had similar expression level in both blastomere types, such as the housekeeping gene *rps-2*. **(Figure 2B-C)**. The identity (somatic versus germline) of the sorted blastomeres was also verified (see **Technical Validation**).

**Figure 1.**
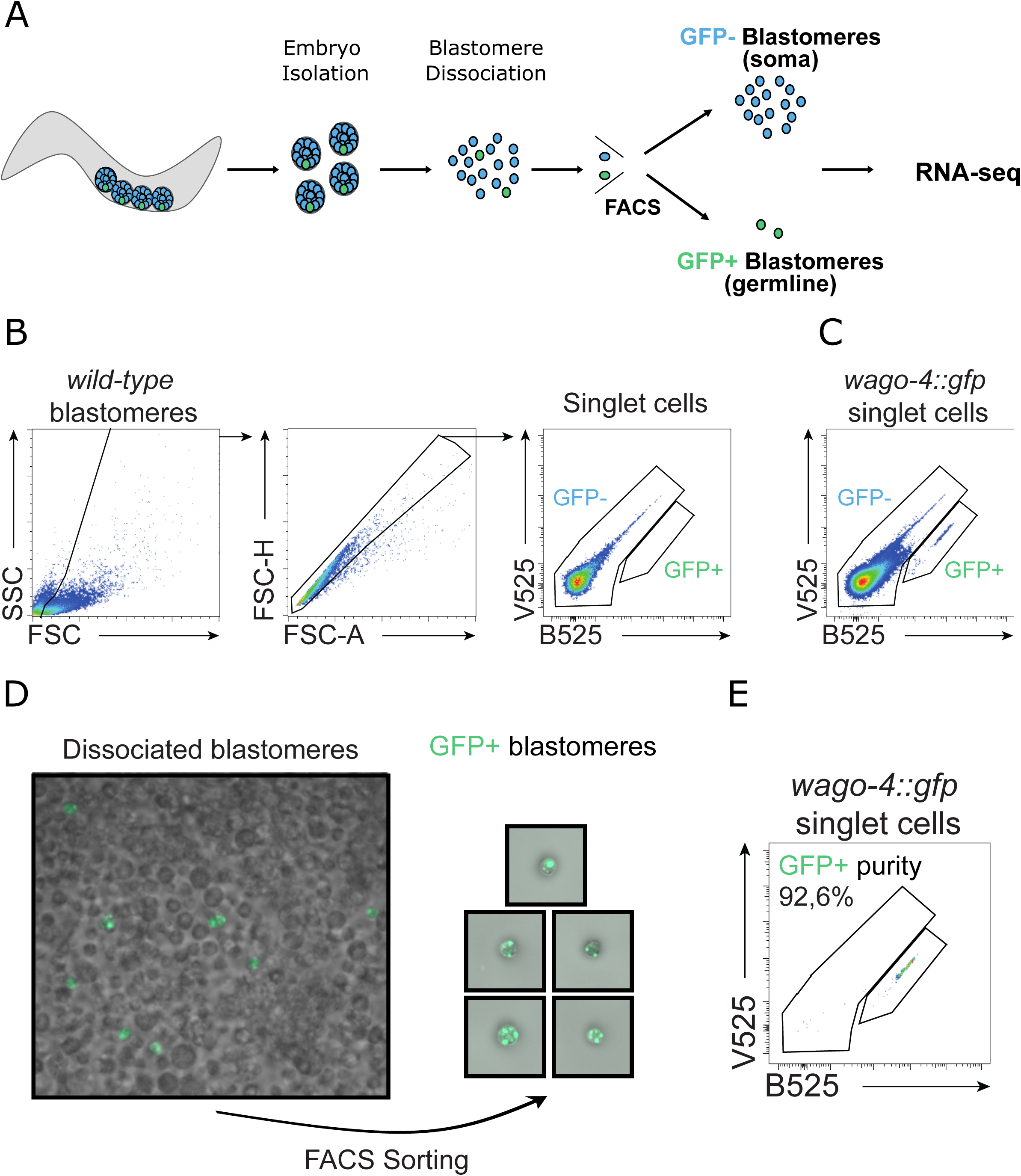
FACS analysis of germline and somatic blastomeres in the *wago-4::gfp* strain. **A.** Schematic illustrating the workflow for RNA-seq performed on GFP+ (germline) separated from GFP- (somatic) blastomeres by FACS. **B.** Gating strategy. Blastomeres were first gated based on the side scatter (SSC) and forward scatter (FSC) to eliminate debris. Subsequently, singlet cells were identified based on forward scatter-height (FSC-H) versus forward scatter-area (FSC-A). Finally, GFP+ blastomeres were identified by excitation at 405 nm (detected with the V525 filter) and 488 nm (detected with the B525 filter). No GFP+ blastomeres were observed in the *wild-type* strain, whereas they were present in the *wago-4::gfp* strain. **C.** Imaging of blastomeres isolated from the *wago-4::gfp* strain before FACS and imaging of the GFP+ blastomeres after FACS. **D.** Purity assessment of the GFP+ blastomeres by flow cytometry.

**Figure 2.**
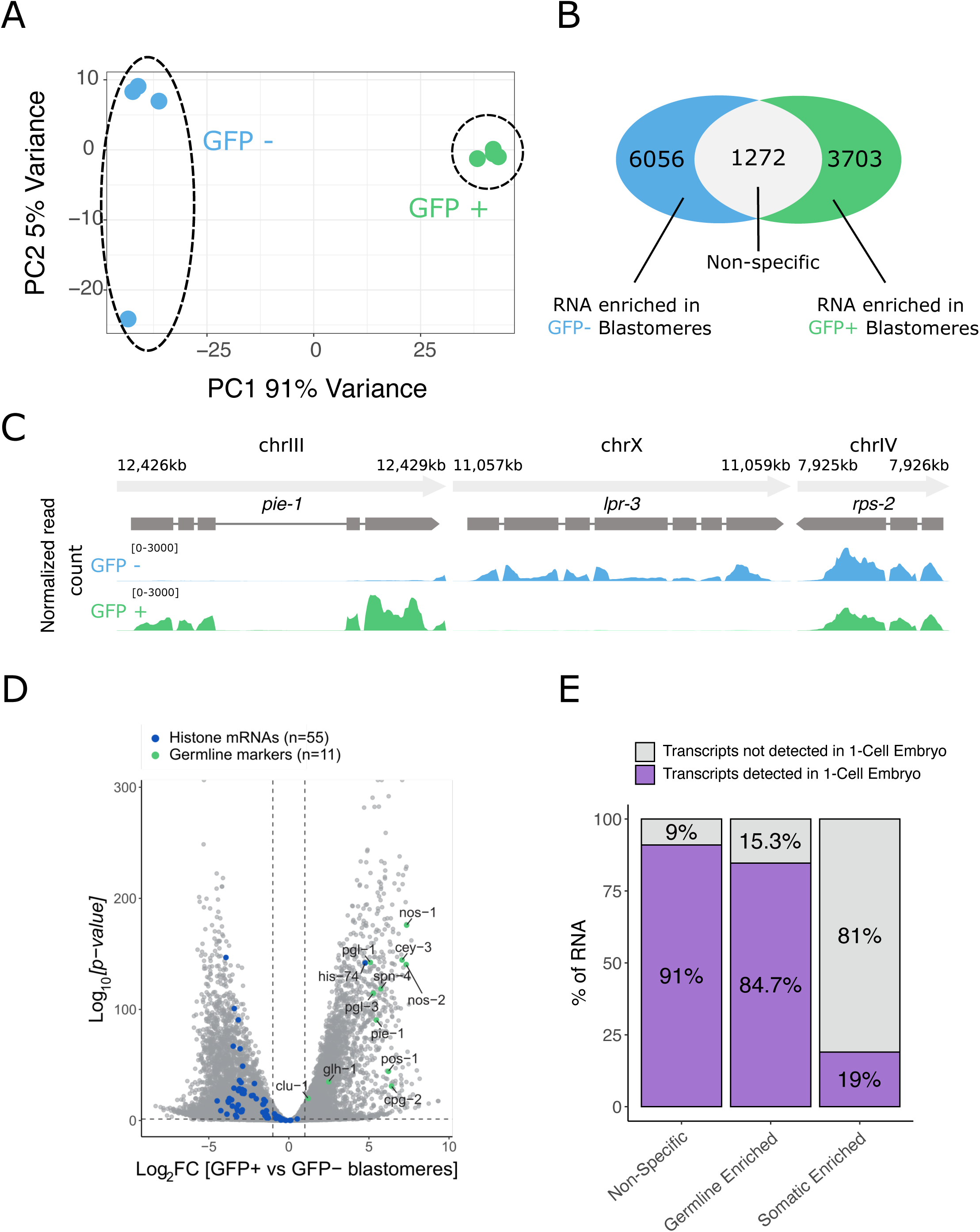
RNA-seq performed on FACS-sorted somatic and germline *C. elegans* blastomeres. **A.** Principal Component Analysis (PCA) of GFP- (blue) and GFP+ (green) blastomeres, with the first two principal components explaining 96% of the variance (n = 4 biological replicates). **B.** Venn diagram showing the number of transcripts enriched in GFP- blastomeres (blue), non-specific (grey) or enriched in GFP+ blastomeres (green). Transcripts with a Fold change of +/- 2 were considered as enriched in GFP- or GFP+ blastomeres. Only transcripts with an adjusted p value < 0.01 (total of 11031) were considered. **C.** Normalized RNA-seq read coverage for one replicate for one RNA enriched in GFP-blastomeres (*lpr-3),* one non-specific RNA (*rps-2)* and one RNA enriched in GFP+ blastomeres (*pie-1)* in GFP- (blue) or GFP+ (green) blastomeres. **D.** Volcano plot showing the differential expression of transcripts in GFP+ compared to GFP-blastomeres. Green dots highlight 11 germline-specific markers from the literature and blue dots highlight 55 (detected histone mRNAs) of the 64 *C. elegans* histone mRNAs. Dotted lines indicate an adjusted p-value of 0.01 and a Fold change of +/- 2. **E.** Percentage of transcripts detected in the 1-cell stage embryo ^7^ (purple) or not (grey) among the three different categories of transcripts.

We conclude that the WAGO-4-based isolation of germline blastomeres revealed the distinct transcriptomic signatures that characterize each blastomere type. From now on, the transcripts enriched in GFP-/GFP+ blastomeres will be referred to as somatic-enriched RNAs and germline-enriched RNAs respectively whereas transcripts similarly detected in both blastomere types (absolute fold change below 2) as non-specific.

In *C. elegans*, a pool of mRNAs is maternally inherited by the offspring, complicating the detection of *de novo* transcription. Using previous 1-cell embryo RNA-seq data ^7^, which can be considered to represent maternally inherited RNA, we examined whether the three categories of transcripts we defined were already detectable in the 1-cell embryo **(Figure 2E)**. For somatic-enriched RNAs, only 19% (n=1149) appeared to be maternally provided, in agreement with *de novo* transcription being activated in somatic blastomeres. In contrast, 85% (n=3136) of the germline-enriched RNAs were already present in the 1-cell embryo, which suggest they are mostly maternally inherited. Nevertheless, 15% (n=567) were not detected in the 1-cell embryo, suggesting that these may originate from unexpected embryonic transcription in the germline blastomere during early embryogenesis. These RNAs mostly include coding RNAs but also noncoding RNAs (n=58) and pseudogenes (n=102) (**Figure 2E**). These data support that the overall view of the germline blastomeres being completely transcriptionally silent may be too restrictive, consistent with recent findings that germline blastomeres chromatin structure resembles the transcriptionally permissive state of somatic blastomeres ^8^.

### PIE-1 maintains germline identity

To further investigate the role of PIE-1 on transcriptome composition in somatic and germline blastomeres, we used the auxin-inducible degron system (AID) ^9^. Efficient depletion of PIE-1 was also verified (see **Technical Validation**). Therefore, we crossed *pie-1::degron::gfp* into the *wago-4::gfp* background. We observed partial synthetic sterility and embryonic lethality occurring even in the absence of 3-IAA (indole-3-acetic acid) and depletion of PIE-1::degron::GFP **(Figure 3A-B)**. However, depleting PIE-1::degron::GFP led to a complete maternal-effect embryonic lethality (**Figure 3C**). Importantly, WAGO-4::GFP localization remained unaffected **(Figure 3D)**, thereby allowing the sorting of germline blastomeres in embryos depleted of PIE-1.

**Figure 3.**
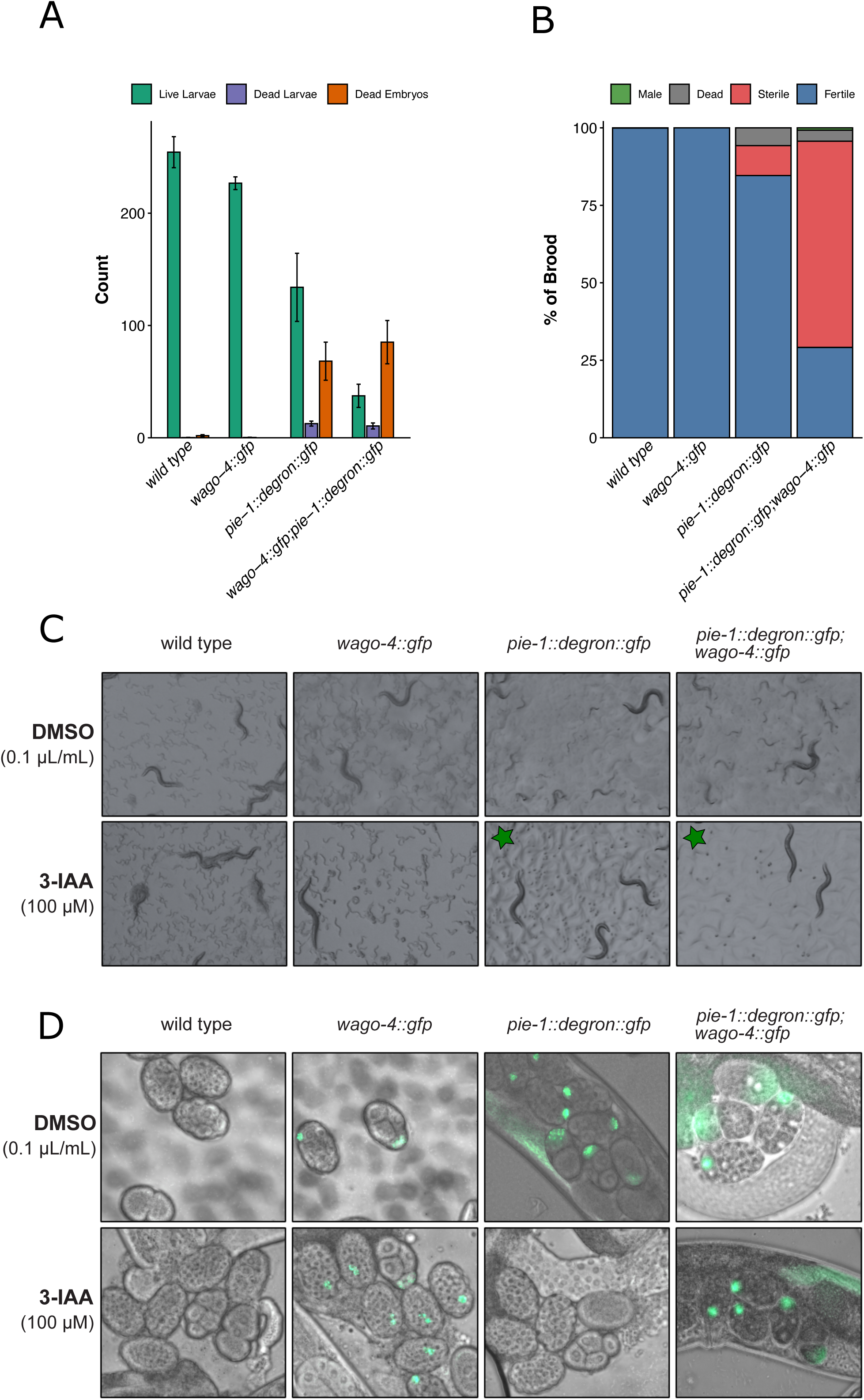
Tagging of WAGO-4 in the *pie-1::degron::gfp* in the presence or absence of 3-IAA) **A.** Number of viable larvae, dead larvae and dead embryos per worm for the indicated strains. Data are presented as means +/- 1 standard deviation for the indicated number of parental worms. **B.** Percentage of the alive larvae in A that became fertile, sterile, male adults or died during the course of development. Data represents the means for the indicated number of parental worms. **C.** Transmitted light images of the indicated strains and conditions. Images were captured 48 to 72 hours after the parental L4 larvae were placed on plates. The adults visible on the plates are the originally deposited parental L4 larvae. Conditions where no F1 larvae (embryonic lethal phenotype) were observed are indicated with a green star. **D.** Fluorescence imaging of the indicated *C. elegans* mutant embryos and conditions. Images were taken 24 hours after the parental L4 larvae were placed on plates. Upon exposure to 3-IAA, the PIE-1::degron::GFP protein was degraded, resulting in the loss of GFP fluorescence in the *pie-1::degron::gfp* strain. However, in the *pie-1::degron::gfp - wago-4::gfp* strain, despite PIE-1::degron::GFP degradation as shown by the embryonic lethality observed in **C**, germline blastomeres could still be accurately identified due to the proper localization of WAGO-4::GFP.

To validate our experimental setup, we first separated germline from somatic blastomeres in the *pie-1::degron::gfp wago-4::gfp* strain without depleting PIE-1. After RNA-seq, a total of 3629 RNAs were identified as specifically enriched in germline blastomeres, 76% of which (n=2776) overlapped with transcripts previously detected in *wago-4::gfp* embryos. These results confirm the successful separation of germline and somatic blastomeres and indicate that the germline transcriptome is consistent across varying strains.

We next repeated RNA-seq on PIE-1 depleted embryos. We observed a marginal effect on somatic blastomere composition **(Figure 4A-B)** with only 161 downregulated, and 13 upregulated transcripts (Fold change +/- 2 and adjusted p-value < 0.01). In contrast, the transcriptome of germline blastomeres underwent significant changes following the depletion of PIE-1 **(Figure 4A),** with 5263 misregulated transcripts (3059 downregulated and 2204 upregulated). We observed (i) a global downregulation of “germline-enriched” transcripts **(Figure 4C**, bottom right), and (ii) an upregulation of “somatic-enriched” transcripts in germline blastomeres **(Figure 4C**, top left). Unexpectedly, 81% (1392/1712) of the downregulated germline-specific RNAs are already detected in the 1-cell embryo **(Figure 4D),** suggesting a new critical role of PIE-1 in maintaining the steady-state of maternally inherited transcripts. In addition, the dataset confirms the previous model of PIE-1 repressing somatic-specific genes. Furthermore, germline-specific markers are no longer enriched in germline blastomeres upon PIE-1 removal **(Figure 4E).**

**Figure 4.**
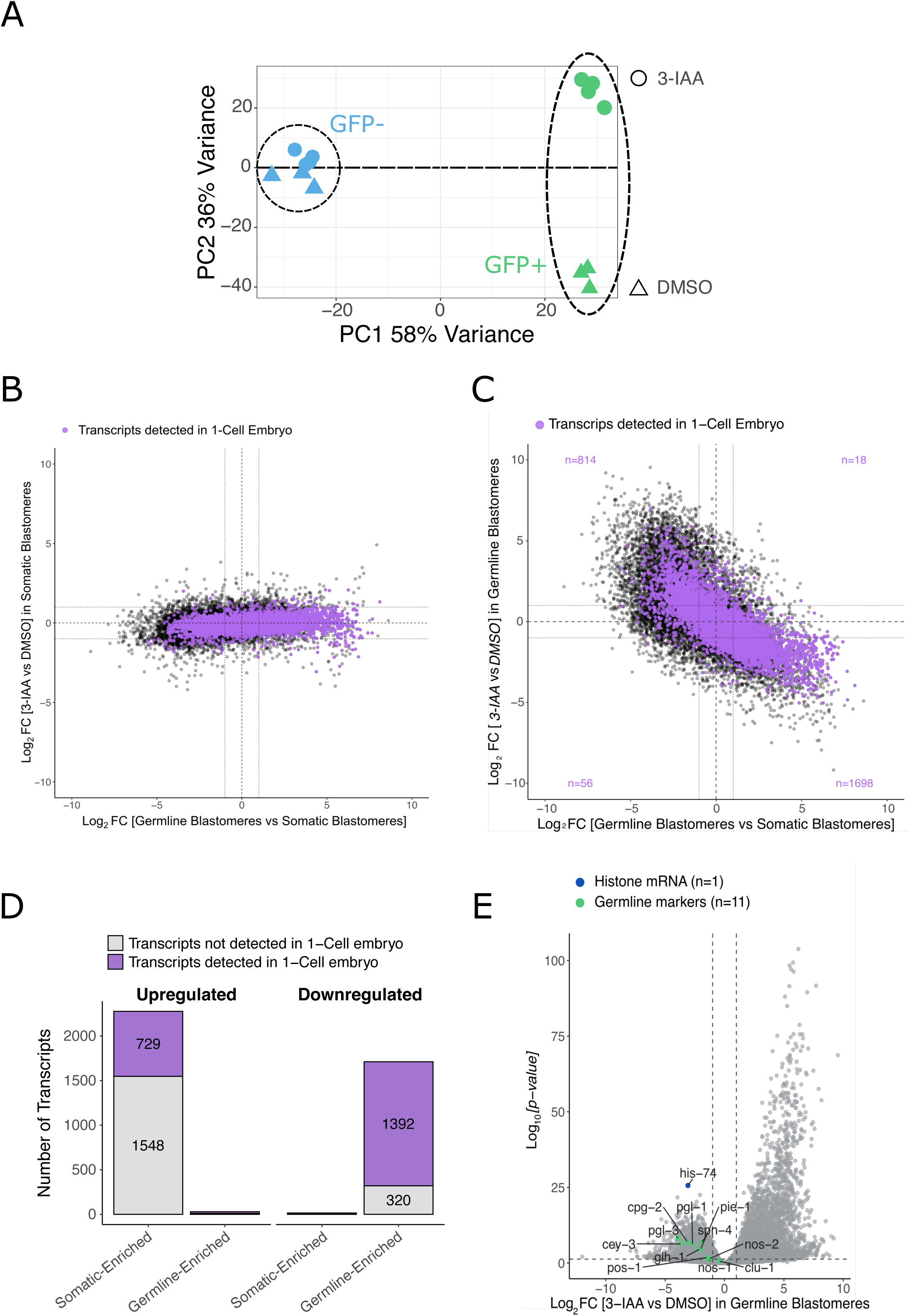
**Effect of PIE-1 depletion on germline blastomeres identity** A. Principal component analysis (PCA), explaining 94% of the variance, showing somatic blastomeres (blue) and germline blastomeres (green) from embryos of L4 larvae treated with DMSO (triangles) or 3-IAA (circles) (n = 3 biological replicates for DMSO treatment and 4 for 3-IAA treatment) B. Differential mRNA expression induced by 3-IAA treatment compared to DMSO in somatic blastomeres (Y-axis) for germline (right) or somatic (left)-enriched RNAs (X-axis). Each transcript with two non-null adjusted *p* values is represented by a dot. Purple dots highlight transcripts detected in the 1-cell stage embryo. The dotted lines indicate a Fold change of +/- 2. C. Differential mRNA expression induced by 3-IAA treatment compared to DMSO in germline blastomeres (Y-axis) for germline (right) or somatic (left)-enriched RNAs (X-axis). Each transcript with two non-null adjusted *p* values is represented by a dot. Purple dots highlight transcripts detected in the 1-cell stage embryo. The dotted lines indicate a Fold change of +/- 2. D. Number of transcripts detected in the 1-cell stage embryo ^7^ (purple) or not (grey) for each quadrant of **C**. Only transcripts with two adjusted p values < 0.01 and two-Fold change of +/- 2 were considered. Upregulated and downregulated represent transcripts that were downregulated or upregulated by the 3-IAA treatment compared to the DMSO in germline blastomeres (Y-axis from B). Germline or somatic-enriched represent transcripts that are germline or somatic-enriched (X-axis from B). E. Volcano plot showing the differential expression of transcripts in germline blastomeres treated with 3-IAA compared to those treated with DMSO. Green dots highlight 11 germline-specific markers from the literature, and the blue dot highlights the germline-specific mRNA *his-74*. Dotted lines indicate an adjusted p-value of 0.01 and a Fold change of +/- 2.

The present dataset reinforces the concept of PIE-1 acting as a molecular safeguard that preserves germline integrity. Indeed, RNA-seq analyses of sorted germline and somatic blastomeres **(Figure 4)** demonstrated that the absence of PIE-1 results in the downregulation of germline specific RNAs. The majority of those RNAs are of maternal origin, which suggests that PIE-1 controls their level independently of their transcription. Moreover, PIE-1 prevents somatic transcripts from accumulating in the germline blastomeres.

### *C. elegans* strains and media

All *C. elegans* strains were derived from the Bristol N2 strain. Worms were maintained at 20°C on NGM-agar plates (Nematode Growth Media – agar: 51.3 mM NaCl, 1 mM CaCl2, 1 mM MgSO4, 25 mM KH2PO4, 5 µg/mL cholesterol and 1.25g peptone with 8.5g agar for 500 mL) seeded with *E. coli* OP50. In liquid medium, embryos hatched in M9 buffer (42.2 mM Na2HPO4, 22 mM KH2PO4, 85.7 mM NaCl, 1 mM MgSO4). A list of *C. elegans* strains used in this thesis is included in the **Supplementary Table S1**.

### C. elegans crossing

Males of the *wago-4::gfp* strain were generated by incubating L4 hermaphrodites at 30°C for 4–6 hours, followed by a return to 20°C to allow progeny development. The resulting males were isolated and 10 of them were seeded with a single *pie-1::degron::gfp* hermaphrodite on an NGM plate. The plate was incubated at 20°C for several days. The presence of numerous males among the progeny suggested successful mating. A subset of worms was then isolated, and single-worm PCR was performed to confirm the presence of the desired alleles. All primers used in this study are listed in **Supplementary Table S2**.

### Single worm PCR

Before PCR, worms were digested in 5 µL of 1X Colorless GoTaq Reaction Buffer (Promega M792A) containing 1 mg/mL proteinase K (Roche Diagnostics 03 115 828 001) first by snap-freezing for 15 minutes and then by incubating 1 hour at 65°C to obtain a worm lysate. The proteinase K was inactivated for 15 minutes at 95°C before the worm lysate containing DNA was used for PCR.

The PCR reaction was performed using GoTaq polymerase (GoTaq G2 DNA Polymerase, Promega M748B) with 1.5 µL of worm lysate from proteinase K treated worms, 5 µL of 5X Green GoTaq Reaction Buffer (Promega M791A), 2 µL of primers at 10 µM, 0.25µL of dNTP at 20 mM, 0.125 µL of GoTaq polymerase and 14.1 µL of water. The annealing temperature corresponded to the Tm (Primer Melting Temperature) of the primer with the lowest Tm, and the extension time is 1 minute per kb.

### Indole-3-acetic acid treatment

L4 larvae were plated on NGM plates supplemented with 100 µM indole-3-acetic acid (3-IAA), dissolved in DMSO (Thermo Fisher Scientific A10556), for 18 hours. Control L4 larvae were placed on NGM plates containing 0.1 µL/mL DMSO. Adult worms were then collected for embryo and then blastomere isolation.

### *C. elegans* brood size and progeny viability assay

On day 0, one parental L4 worm of each genotype was transferred to an individual plate. On day 1, the worms reached adulthood and were allowed to lay eggs. On day 2, they were transferred to a new plate, and on day 3, they were moved again to a new plate. The first plate was counted on day 3, the second on day 4, and the third on day 5. On each counting day, viable larvae, dead larvae, and unhatched (dead) embryos were recorded. The sum of viable and dead larvae plus unhatched embryos on all 3 plates for a single worm is its total brood size. The ratio of dead eggs to the total brood size is the embryonic mortality.

After counting, the plates were kept until the F1 alive larvae reached adulthood, at which point the number of fertile, sterile, dead or male adults was counted.

### *C. elegans* embryo imaging

Gravid adult worms were killed in 50 mM NaN3 directly on slides and then cut in half with a 22G needle (BD Microlance 3) to release the embryos from the adult body or to adjust their positioning within the body. A coverslip was then placed on each slide, which was then imaged using the Zeiss Axio Observer, outfitted with the Zeiss Colibri 7 Type R^10^CBV-UV system and a Hamamatsu Digital Camera C11440, with image acquisition performed via Zeiss ZEN software.

### *C. elegans* blastomere isolation, FACS and RNA sequencing

L4 larvae of the *wago-4::gfp* strain were plated on NGM plates, while L4 larvae of the *pie-1::degron::gfp wago-4::gfp* strain were plated on NGM plates supplemented with 3-IAA or DMSO for 18 hours, until reaching the gravid adult stage. Embryos (3.000.000 for the *wago-4::gfp* strain and the maximum for the *pie-1::degron::gfp wago-4::gfp* strain) were then isolated from adults via hypochlorite treatment. The lysis reaction was halted after 5 minutes by adding twice the volume of egg buffer (118 mM NaCl, 48 mM KCl, 2 mM CaCl₂, 2 mM MgCl₂, 25 mM HEPES pH 7.3).

Intact embryos were separated from debris by centrifugation at 450 g for 5 minutes in 30% sucrose egg buffer. Eggshells were then removed by incubation in 500 µL of chitinase 1 U/mL (Merck C6137, resuspended in egg buffer) for approximately 20 minutes at RT. When 80% of embryos were eggshell-free, the reaction was stopped by the addition of 1 mL of L15 medium (Gibco 11415-064), and embryos were pelleted at 1300 g for 2 minutes at RT. The pellet was resuspended in 800 µL of L15 medium, and embryos were dissociated by passing the suspension through a 3 mL syringe (Henke-Ject 8300005762) fitted with an 18G needle (Agani needle AN*1838R1). Dissociation was halted when approximately 80% of embryos were dissociated. The suspension was then directly filtered on ice through a 15 µm PET mesh (pluriStrainer 43-50015-50) using 10 mL of cold L15 medium. The filtered cells were then pelleted at 900 g for 3 minutes at 4°C, washed with cold egg buffer containing 1% BSA and 40 U/mL RNase inhibitor (Merck Protector RNase Inhibitor 03335402001), and resuspended in cold egg buffer with 1% BSA and 40 U/mL RNase inhibitor. Cells were kept on ice and immediately processed.

Cell sorting was conducted on a Beckman Coulter Cytoflex SRT with the single-cell sorting mask. Forward scatter (FSC) and side scatter (SSC) were used for gating, and singlet cells were identified based on FSC-H versus FSC-A. To distinguish real GFP signal from autofluorescence, two lasers were employed: 405 nm and 488 nm, as GFP is also excited by the violet laser. GFP fluorescence was detected using 525/40 BP filters on each laser, called V525 and B525. Cells were sorted at a flow rate of 40 µL/min directly into Eppendorf tubes containing 1 mL of Trizol LS (Invitrogen 10296010). Between 13,000 and 50,000 cells were sorted depending on the replicate and, for each replicate, equal numbers of GFP-positive and GFP-negative cells were collected. A *wild-type* strain was used as a control to appropriately gate GFP-positive cells.

100 µL of 1-bromo-3-chloropropane (Merck B9673) was then added to the sorted cells in Trizol LS, and the sample was incubated at RT for 15 minutes. It was then centrifuged at 12000 g for 15 minutes. The upper aqueous phase was collected and combined with 2 µL of GlycoBlue coprecipitant (Invitrogen AM9515) and 500 µL of isopropanol, followed by another 15-minute incubation at RT. RNA was pelleted by centrifugation at 12000 g for 15 minutes, washed with 75% ethanol, air-dried, and resuspended in DEPC-treated (diethyl pyrocarbonate) water. Genomic DNA was then removed using Turbo DNase (Thermo Fisher Scientific AM2238) according to the manufacturer’s protocol. RNA was further purified by phenol-chloroform extraction. The volume of RNA was adjusted to 400 µL with DEPC water, and 40 µL of 3 M sodium acetate (pH 5.2) and 880 µL of phenol-chloroform-isoamyl alcohol (125:24:1) (Merck 77619) were added. The sample was centrifuged at 14,000 g for 2 minutes. The upper phase was collected, mixed with twice the volume of chloroform-isoamyl alcohol (24:1) (Merck 25666) and centrifuged again. The upper phase was recovered, and 1.5 µL of GlycoBlue coprecipitant along with 2.5 times the volume of 100% ethanol was added. The RNA was precipitated at -20°C for 1 hour, pelleted at 12000 g for 30 minutes, washed with 75% ethanol, air-dried and resuspended in DEPC-treated water.

RNA concentration and integrity were assessed on a TapeStation (Agilent) using the High Sensitivity RNA ScrenTape kit (5067-5579). Ribosomal and mitochondrial RNAs were removed using a custom depletion strategy (Seqalis). RNA-seq library preparation was made using the Illumina Total RNA prep with Ribo-Zero Plus (20040529) and the Twist Mechanical Fragmentation Library Preparation Kit (101281) (Seqalis).

Reads were mapped to the *C. elegans* genome (version WBcel235.101) with HISAT2^11^ with options --no-mixed --no-discordant --min-intronlen 20 –max-intronlen 5000. Gene-level read quantification was made with featureCounts ^12^ with options --countReadPairs -M -p -T 6. Differential gene expression between sorted cells was made on three (3-IAA vs DMSO) or four (Soma vs Germline) replicates using DEseq2 ^13^. Significant differentially expressed genes were genes defined as genes with an absolute log2 Fold change > 1 and an adjusted *p* value (false discovery rate) < 0.01.

### Data Record

Stubbe, F-X. PIE-1 plays a dual role in defining the C. elegans germline blastomere transcriptional identity during early embryogenesis (RNAseq - P cells). *NCBI Gene Expression Omnibus* https://identifiers.org/geo:GSE309369 (2028).

Stubbe, F-X. PIE-1 plays a dual role in defining the C. elegans germline blastomere transcriptional identity during early embryogenesis (RNAseq – Degron). *NCBI Gene Expression Omnibus* https://identifiers.org/geo:GSE309483 (2028).

### Technical Validation

To verify the identity of the sorted blastomeres, we analyzed a set of 11 known germline-specific markers ^7,14–16^. We observed a strong enrichment of these markers in the GFP+ blastomeres **(Figure 2D**), validating the GFP+ blastomeres as germline blastomeres. In addition, replication-dependent histone mRNAs exhibited lower expression in GFP+ blastomeres, except for the germline-specific *his-74* mRNA ^17^, which is in line with the slower division rate of germline blastomeres and the cell cycle arrest of Z2 and Z3 **(Figure 2D)**.

To verify the efficiency of PIE-1::degron::GFP depletion, we show that it recapitulates the penetrant maternal-effect embryonic lethality of *pie-*1 null alleles **(Figure 3)**.

## Data availability

The transcriptomic datasets generated during the current study are available in the Gene Expression Omnibus (GEO) repository. The RNA-seq data for the *C. elegans wago-4::gfp* strain can be accessed under accession number **GSE309369**. The RNA-seq data for the *pie-1::degron::gfp;wago-4::gfp* strain are available under accession number **GSE309483**.

## Code Availability

The data analysis was performed using HISAT2 and DEseq2. Graph rendering was performed in R using the Tidyverse packages. No custom code or proprietary scripts were developed for this study.

## Acknowledgements

We thank Céline Legrand and Kevin Willemart (UNamur MORPH-IM platform) for support with flow cytometry, Violette Buffet and Germain Agazzi for assistance with statistical computing and graphics on “R”, Carine Michiels and Severin Ronneau for comments on the manuscript.

Pauline Ponsard was a FNRS Research fellow. Damien Hermand worked at the University of Namur until December 2023 and is an honorary FNRS Director of Research.

## Authors contribution

P.P. and D.H. designed the project. P.P. performed the experiments. F.X.S. analyzed the RNA-seq data. P.T. helped with maintaining worms. C.C.M. provided scientific guidance. D.H. acquired the fundings and supervised the project. P.P., F.X.S. and D.H. wrote the paper with inputs from all authors.

## SUPPLEMENTARY TABLES

**Supplementary Table S1:**
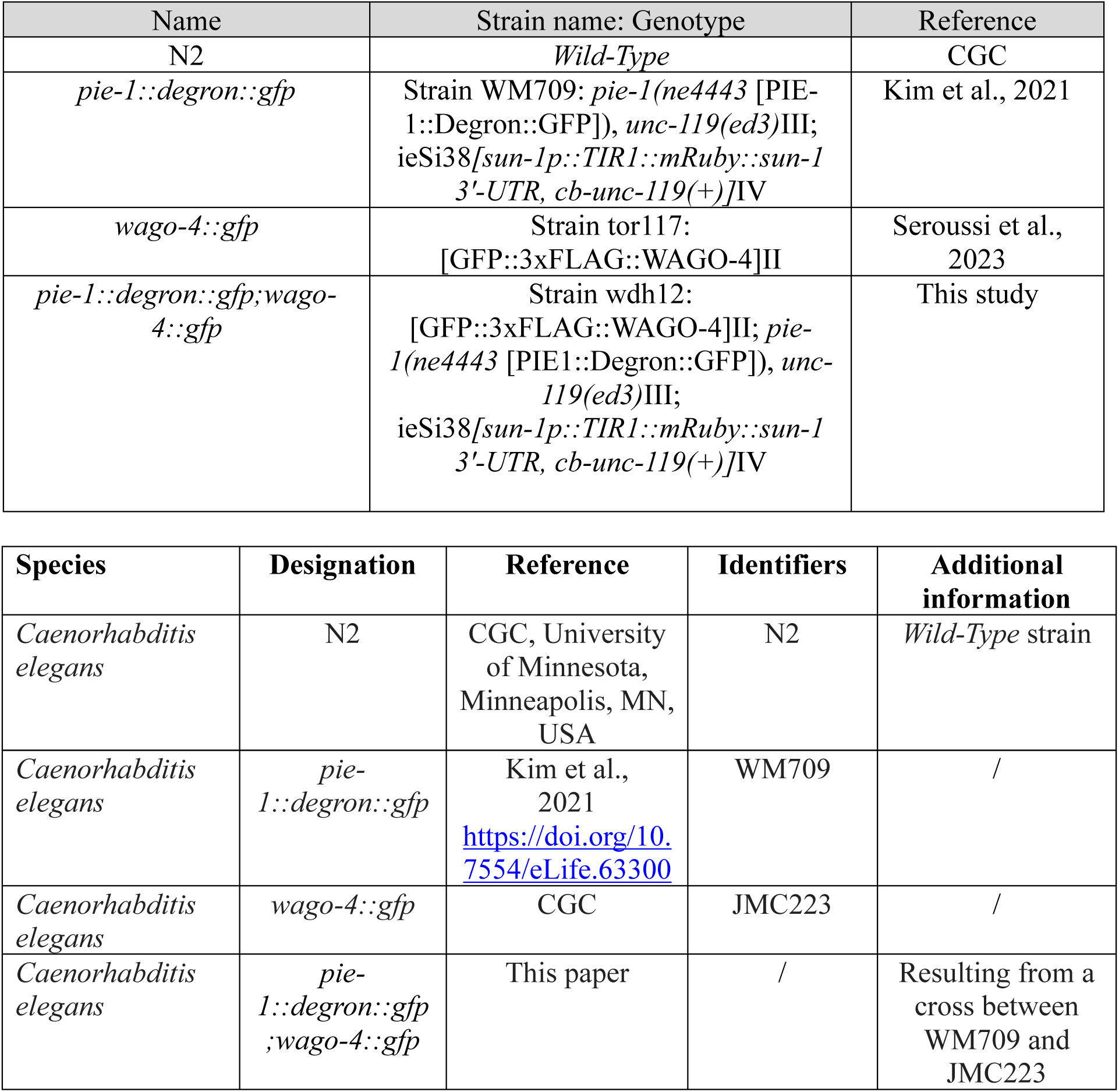
*Caenorhabditis elegans* strains used in this study.

**Supplementary Table S2:**
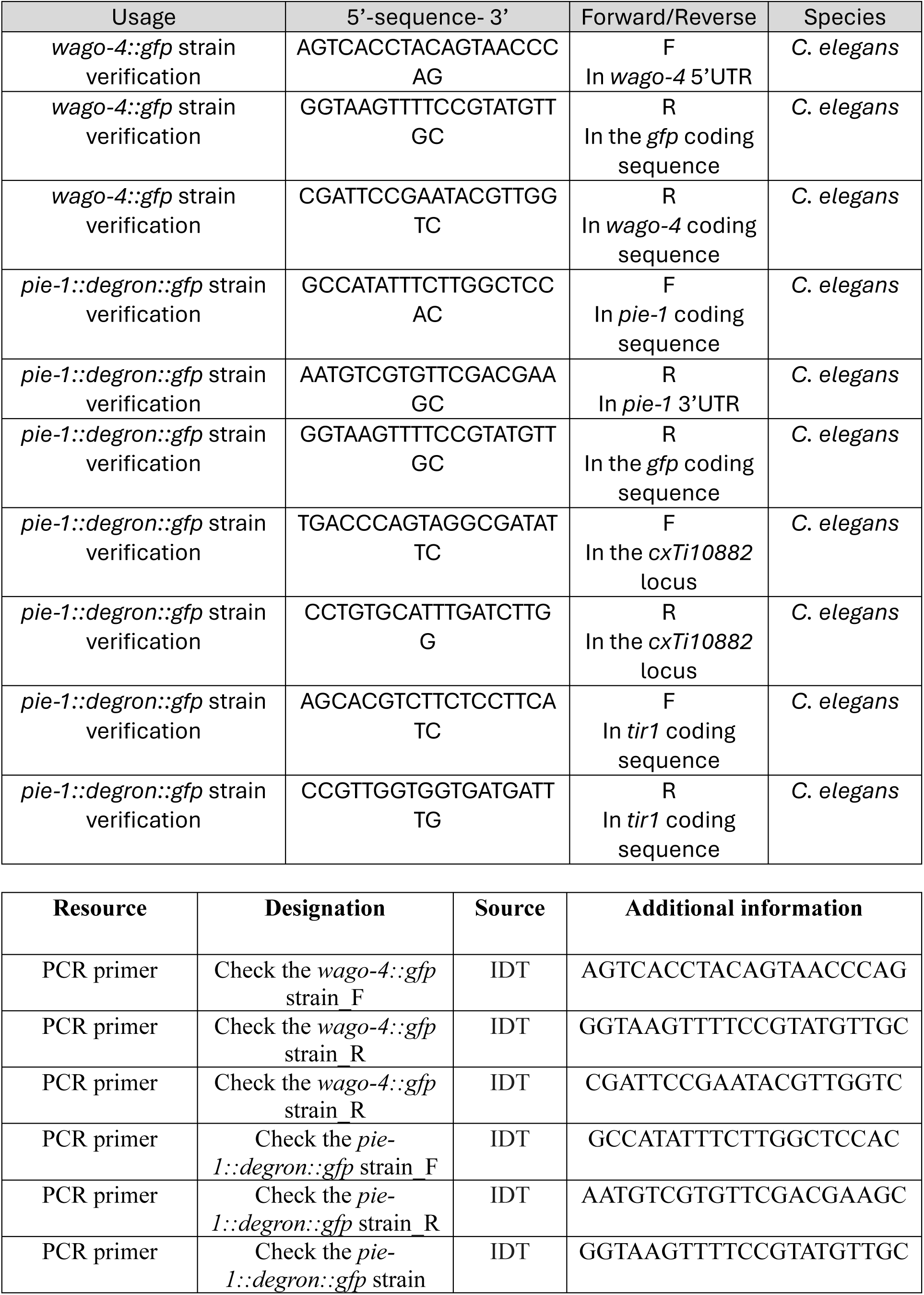
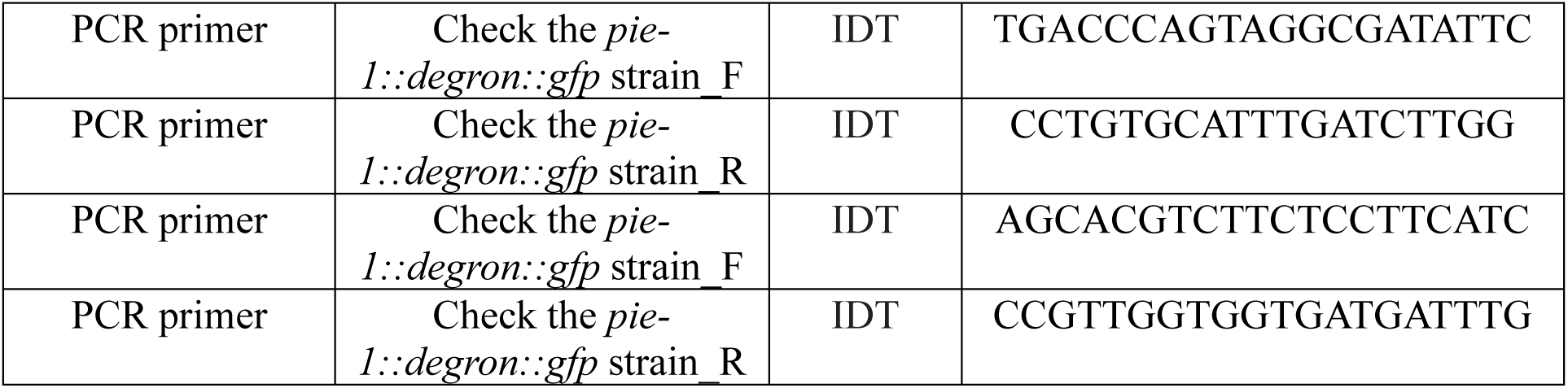
Primers used in this study.

